# Chromosome-scale Genome Assembly of the Allopolyploid *Arundo donax*

**DOI:** 10.1101/2023.06.18.544523

**Authors:** Mengmeng Ren, Fupeng Liu, Xiaohong Han, Daohong Wu, Hai Peng

## Abstract

*Arundo donax* L (*A. donax*). is a promising energy crop with high biomass and wide adaptability, while lack of reference genome limiting the genetic improvement of this crop. Here, we report two different chromosome-scale assemblies of *A. donax* genome (0004 and 0408) using Pacbio SMRT sequencing and Hi-C technology. The genome size of these two assemblies is 1.30 Gb / 2.86 Gb with contig N50 33.15 Mb / 34.62 Mb respectively. A total of 74,403 / 167,586 gene models were predicted in 0004 and 0408 genome, of which over 90% of genes were functionally annotated. Comparative genome analysis revealed that 0004 is alloenneaploid and 0408 is allohexaploid. Further analysis revealed that *A. donax* undergone strong gene family expansion and two whole-genome duplication events during evolution. Our results will enhance genetic understanding and promote the genetic improvement of *A. donax*.

## 1. Introduction

*Arundo donax* L. (*A. donax*) is a tall perennial grass commonly known as giant reed. *A. donax* is a widely utilized plant species with various applications in bioenergy, papermaking, and even as an ornamental lawn grass due to its fast growth and high biomass production [1–4]. One of the significant characteristics of *A. donax* is its high productivity and fast growth rate. The plant matures rapidly and can produce large quantities of biomass in a short time, making it an attractive crop for bioenergy production [5–7]. Additionally, *A. donax* is highly tolerant of a broad range of environmental conditions, including drought, excess water, and high salinity [8–9]. These attributes make the plant an excellent candidate for sustainable agriculture and bioenergy development.

*A. donax* is also known for its environmental and ecological benefits. It is considered to be an excellent tool for phytoremediation, which could be used to clean up contaminated soil and water [10–11]. *A. donax* is effective in removing heavy metals and other pollutants from contaminated sites, making it a valuable resource in environmental restoration [12]. In recent years, there has been increased interest in the study and utilization of *A. donax*, which has led to a number of research efforts aimed at understanding its biological properties.

The sterile nature limits the genetic variability and delays the genetic improvement of *A. donax* [13–16]. However, the fundamental research of *A. donax* genome encountered great difficulties. The ploidy of *A. donax* is still unclear because of the high chromosome number and small size of each chromosome [17–18]. Similar to another aquatic plant *Phragmites australis* [19], *A. donax* showed elastic chromosome number with variable ploidy. Different chromosome number of *A. donax* were reported by several authors, including 110 [20–21], 108 [22], and 84 chromosomes [23]. The complexity of *A. donax* genome hinders further understanding and the genetic improvement of this valuable energy crop.

The large and complex nature of plant genomes, coupled with varying degrees of ploidy, has long posed challenges for genome assembly in plants. Fortunately, advancements in sequencing technologies, such as third-generation long-read sequencing, particularly High Fidelity (HiFi) reads sequencing, along with new assembly algorithms serve as the foundation for the rapid advancements in plant genome sequencing [24]. HiFi sequencing allows for the generation of long-read, high-quality data that can be used to accurately assemble and annotate complex genomes. It has already proven to be a valuable tool in decoding complex plant genomes, such as allohexaploid oat [25], tetraploid potato [26], and autotetraploid alfalfa [27].

Here, we assembled the first two chromosome-scale *A. donax* genome by combining Pacbio SMRT sequencing and high-throughput chromosome conformation capture (Hi-C) technology. These two assemblies will facilitate the better understanding of *A. donax* genomics and better utilization of *A. donax* resources.

## 2. Materials and Methods

### 2.1. Plant materials

The two *A. donax* plants were collected from Wuhan, Hubei (0004) and Yueyang, Hunan (0408), respectively. About 3 cm stem with a tiller bud were used as explant to propagate with tissue culture. Seedlings were then acclimatized outside and transplanted in the field.

### 2.2. Karyotype analysis

Karyotype analysis was conducted followed a previously report [28–29]. Briefly, root tips about 2 cm in length were placed in Nitrous Oxide (0.9-1.0 Mpa) for two hours, and fixed with pre-cold glacial acetic acid. After that, Root tips were digested with cellulase and pectinase at 37°C for 1 h and crushed using a dissecting needle. Cells suspension were observed under an optical microscope to get the suitable chromosome specimens. Fluorescent *in situ* hybridization probes were prepared using the nick translation method. Probes for telomere, 5SrDNA and 18SrDNA are fluorescently-labeled (TTTAGGG)6 oligonucleotides, pTa794 and pTa71 plasmids respectively. After hybridization, slides were photographed under a fluorescence microscope.

### 2.3. Flow cytometry analysis

Karyotype analysis was conducted followed a previously report [30]. Samples were immersed into MGb dissociation solution (45 mM MgCl_2_·6H_2_O, 20 mM MOPS, 30 mM sodium citrate, 1% (W/V) PVP, 0.2% (v/v) Tritonx-100, 10 mM Na_2_EDTA, 20 μL/mL β-mercaptoethanol pH 7.5), chopped with blade and filtered to get the nuclear suspension. The suspension was then incubated in propidium iodide and RNase for 1 h and detected with flow cytometry (BD FACScalibur). Tomato and barley were used as internal references.

### 2.4. DNA extraction, Pacbio library preparation and sequencing

High molecular weight genomic DNA was prepared with QIAGEN® Genomic kit (13343) according to the manufacturer’s instructions. SMRTbell libraries constructed PacBio’s standard protocol. Briefly, the quality qualified DNA was sheared by g-TUBEs (Covaris, USA), damage repaired and 20-50 kb DNA fragments were screened by the BluePippin (Sage Science, USA) and purified using AMPure PB beads (Agilent technologies, USA). Sequencing was performed on a PacBio Sequel instrument with Sequel II Sequencing Kit 2.0. Raw sequencing data were filtered through SMRTlink 8.0.

### 2.5. MGISEQ-2000 library preparation and sequencing

Genomic DNA was extracted using the CTAB method and subjected to sonication by Covaris. The DNA fragments range from 200-400 bp were selected using Agencourt AMPure XP-Medium kit. After end-repair, 3’ adenylated, adapters-ligation and PCR Amplifying, the libraries were prepared using Hieff NGS® Fast-Pace DNA Cyclization Kit (Yeasen, 13341ES96) and sequencing on MGISEQ-2000 platform.

To facilitate the annotation of the genome, we performed mRNA sequencing on four plant tissues including leaf, root, panicle and lateral bud. We purified up to 400 µg of total RNA per sample using the TRIzol-based method (Invitrogen, USA), and subsequently treated it with DNase I. PolyA mRNA was enriched using the protocol of NEBNext ® Poly(A) mRNA Magnetic Isolation Module (New England Biolabs #7490 S, USA). RNA sequencing libraries were then prepared using Hieff NGS® Ultima Dual-mode mRNA Library Prep Kit (Yeasen, 12309ES96) following the manufacturer’s instructions. The RNA library concentration was determined using the Qubit ® 3.0 Fluorometer, and the libraries were then subjected to 150 bp paired-end sequencing on a MGISEQ-2000 platform. In total, we generated 55.8 Gb / 51.9 Gb raw data for the four tissue samples of 0004 and 0408, respectively.

### 2.6. Estimation of genome size and heterozygosity

The genome size of *A. donax* was estimated based on k-mer analysis using KMC3 program [31]. The GCE and FindGSE software were used to analyze the 17-mer frequency distribution and perform genome size estimation [32]. We estimated the genome heterozygosity by combining simulated data with different heterozygosity in the Arabidopsis genome and the distribution of 17 k-mer data of *A. donax*.

### 2.7. De novo assembly

The quality-filtered HiFi reads were used for genome assembly with hifiasm (v.0.12) [33] to obtain the preliminary assembly. The completeness of genome was assessed using BUSCO (v.4.0.5) [34] and CEGMA (v.2) [35]. To assess the accuracy of the assembly, all the MGISEQ-2000 reads were mapped to the genome using BWA [36], and SAMtools [37] and bcftools [38] were used to calculate the error rate of genome assembly. Besides, LTR Assembly Index (LAI) was calculated to assess the assembly quality [39].

### 2.8. Hi-C scaffolding

To attach the scaffolds to chromosomes, we extracted genomic DNA using young leaves for the Hi-C library, and the sequencing data was obtained via the MGISEQ-2000 platform. To elaborate, we first cut freshly harvested leaves into 2 cm pieces and vacuum infiltrated them with nuclei isolation buffer supplemented with 2% formaldehyde. Crosslinking was stopped by adding glycine and performing additional vacuum infiltration. We then converted the fixed tissue into powder and re-suspended it in nuclei isolation buffer to obtain a nuclei suspension. *Dpn*II was used to digest the purified nuclei with 100 units, and after marking with biotin-14-dCTP, biotin-14-dCTP from non-ligated DNA ends was removed through the exonuclease activity of T4 DNA polymerase. The ligated DNA was sheared into 300-600 bp fragments, followed by blunt-end repair and A-tailing before purification through biotin-streptavidin-mediated pull down. Finally, the Hi-C libraries were quantified and sequenced using the MGISEQ-2000 platform.

In total, 375 Gb and 347 Gb paired-end reads were generated for 0004 and 0408. To ensure the quality of the Hi-C raw data, we employed HiC-Pro (v2.8.1), a previously established quality control tool [40]. We first filtered out low-quality sequences (quality scores < 20), adapter sequences, and sequences shorter than 30 bp using fastp [41]. The clean paired-end reads were then mapped to the draft assembled sequence using bowtie2 (v2.3.2) with the parameters “-end-to-end --very-sensitive -L 30” to obtain the unique mapped paired-end reads [42]. We identified and retained valid interaction paired reads using HiC-Pro from the unique mapped paired-end reads for further analysis. Additionally, we filtered out invalid read pairs, including dangling-end, self-cycle, re-ligation, and dumped products, using HiC-Pro. We further clustered, ordered, and oriented the scaffolds onto chromosomes using LACHESIS [43], with parameters CLUSTER_MIN_RE_SITES = 100, CLUSTER_MAX_LINK_DENSITY = 2.5, CLUSTER NONINFORMATIVE RATIO = 1.4, ORDER MIN N RES IN TRUNK = 60, ORDER MIN N RES IN SHREDS = 60. Finally, we manually adjusted any placement and orientation errors exhibiting obvious discrete chromatin interaction patterns.

### 2.9. Repeat element Annotation

To identify tandem repeats and transposable elements in the genome, we used GMATA [44] and Tandem Repeats Finder (TRF) software [45]. GMATA identified simple repeat sequences (SSRs), while TRF recognized all tandem repeat elements in the genome. To identify transposable elements, we used a combination of ab *initio* and homology-based methods. An *ab initio* repeat library for the genome was predicted using MITE-hunter [46] and RepeatModeler with default parameters, including LTR_FINDER [47], LTRharvest [48], and LTR_retriever [49] for plant genome. The resulting library was aligned to TEclass Repbase (http://www.girinst.org/repbase) to classify each repeat family’s type. To identify repeats throughout the genome, we applied RepeatMasker [50] to search for known and novel TEs using the *de novo* repeat library and Repbase TE library. Finally, we collated and combined overlapping transposable elements belonging to the same repeat class.

### 2.10. Gene Prediction

To predict genes in a repeat-masked genome, we employed three distinct approaches: ab initio prediction, homology search, and reference-guided transcriptome assembly. Homolog prediction was performed using GeMoMa by aligning homologous peptides from related species to the assembly to obtain gene structure information [51]. For RNAseq-based gene prediction, we aligned filtered MGISEQ-2000 reads to the reference genome using STAR [52] (default settings) and then assembled the resulting transcripts with stringtie2 [53]. Open reading frames (ORFs) were predicted using PASA [54]. Ab initio gene prediction was performed with Augustus [55], utilizing the default parameters with the training set. We then employed EVidenceModeler (EVM) [56] to produce an integrated gene set, removing genes with TE using the TransposonPSI package (http://transposonpsi.sourceforge.net/) and filtering miscoded genes. Untranslated regions (UTRs) and alternative splicing regions were identified using PASA based on RNA-seq assemblies. We retained the longest transcripts for each locus, while regions outside the ORFs were designated UTRs.

### 2.11. Functional annotation of gene models

Gene function information, as well as motifs and domains of their proteins, were assigned by comparing them with various publicly available databases, including SwissProt, NR, KEGG, KOG, and Gene Ontology. We utilized the InterProScan program with default parameters to identify the putative domains and GO terms of the genes [57]. We also employed BLASTp to compare the EvidenceModeler-integrated protein sequences against four well-known public protein databases, using an E-value cutoff of 1e-05. We selected the results with the lowest E-value. Results from the searches of these five databases were then concatenated to obtain a comprehensive gene function annotation.

### 2.12. Annotation of non-coding RNAs (ncRNAs)

For the identification of ncRNA, we utilized two main strategies: searching against databases and prediction with models. To predict transfer RNAs (tRNAs), we applied tRNAscan-SE was applied with eukaryote parameters [58], andInfernal cmscan [59] was employed to search the Rfam database for microRNA, rRNA, small nuclear RNA, and small nucleolar RNA detection. Additionally, we utilized RNAmmer to predict the rRNAs and their subunits [60].

### 2.13. Comparative genomic analysis

Pairwise alignment between 0004 and 0408 genome were performed using Minimap2 software [61] with parameter: -cx asm10. The paf file were used to obtain the dotplot using R package dotPlotly (https://github.com/tpoorten/dotPlotly/). Collinearity analysis were carried out using MCScan (Python version) software. Dotplots for genome pairwise synteny was visualized with the command ‘python -m jcvi.graphics.dotplot’.

To investigate the evolutionary history of *A. donax*, five grass families, *Oryza sativa*, *Zea mays*, *Sorghum bicolor*, *Setaria italica*, *Saccharum spontaneum* and one dicot plant *Arabidopsis thaliana* along with 0004 and 0408 were used for orthologous analysis, phylogenetic analysis and gene family expansion and contraction analysis using OrthoVeen3 [62] with default parameters.

Whole-genome duplication analysis were performed using wgdi [63]. The synonymous substitution values (Ks) of syntenic gene pairs were calculated with default paramiters and visualized using R program.

## 3. Results

### 3.1. Phenotypic characteristic and cytological analysis of A. donax

In the process of collecting domestic *A. donax* germplasm resources, we identified two different types of *A. donax* in China. Based on the leaf color, we arbitrarily name them common and emerald *A. donax*. Obvious differences can be observed between common and emerald *A. donax*. For example, common *A. donax* shows higher and looser architecture than emerald *A. donax* (Figure 1A). The panicle of common *A. donax* is larger and tighter than that of emerald *A. donax* (Figure 1B). The leaf common *A. donax* is light-green, while emerald *A. donax* is dark-green (emerald) (Figure 1C). The rhizome of common *A. donax* is larger than that of emerald *A. donax* (Figure 1D). Brown hair can be observed on sheath of shoot in common *A. donax*, but not in emerald *A. donax* (Figure 1E). Therefore, to fully dissect the genome of *A. donax*, we selected a common variety 0004 and an emerald variety 0408 for genome assembly.

**Figure 1.**
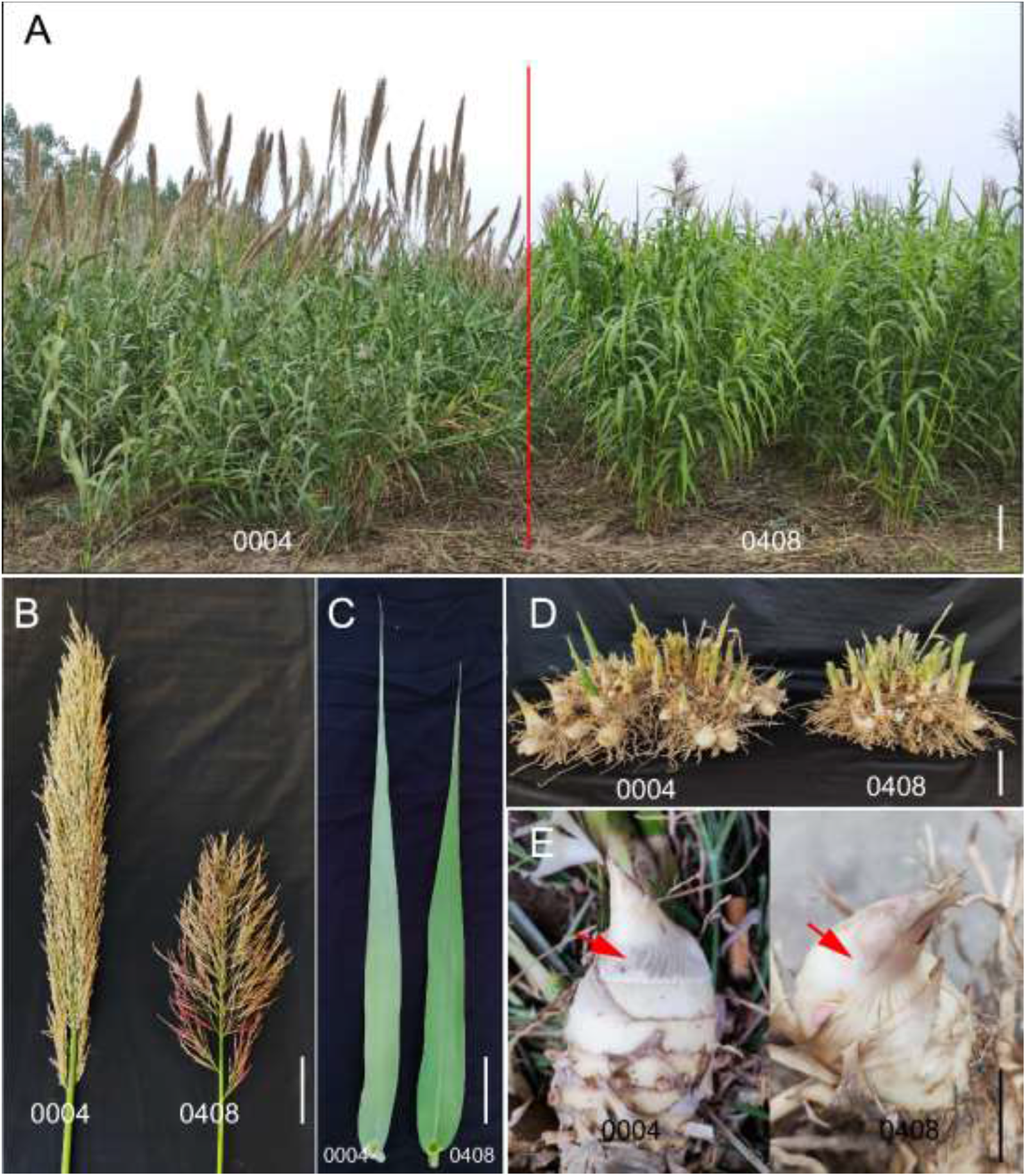
Morphology characteristics of the common and emerald *A. donax*. (A) Plant architecture. (B) Panicle architecture. (C) Leaf morphology. (D) Rhizome morphology. (E) Sheath of shoot. Bars, (A) 50 cm, (B-E) 5 cm.

We first analyzed the karyotype of the two varieties. DAPI staining *in situ* hybridization of telomere repeat sequences showed that 0004 had 112 (Figure 2A-B) chromosomes. In contrast 0408 had 62 chromosomes (Figure 2D-E), in consistent with the fact that chromosome number in *A. donax* is variable. Moreover, rDNA fluorescence *in situ* hybridization revealed that 6/6 chromosomes in 0004 showed strong hybridization signals of 5S rDNA and 18S rDNA (Figure 2C). In contrast, only 2/2 chromosomes in 0408 showed hybridization signals of 5S rDNA and 18S rDNA (one strong signal and one weak signal) (Figure 2F). These results indicated that 0004 and 0008 may have different chromosome basic number and ploidy.

**Figure 2.**
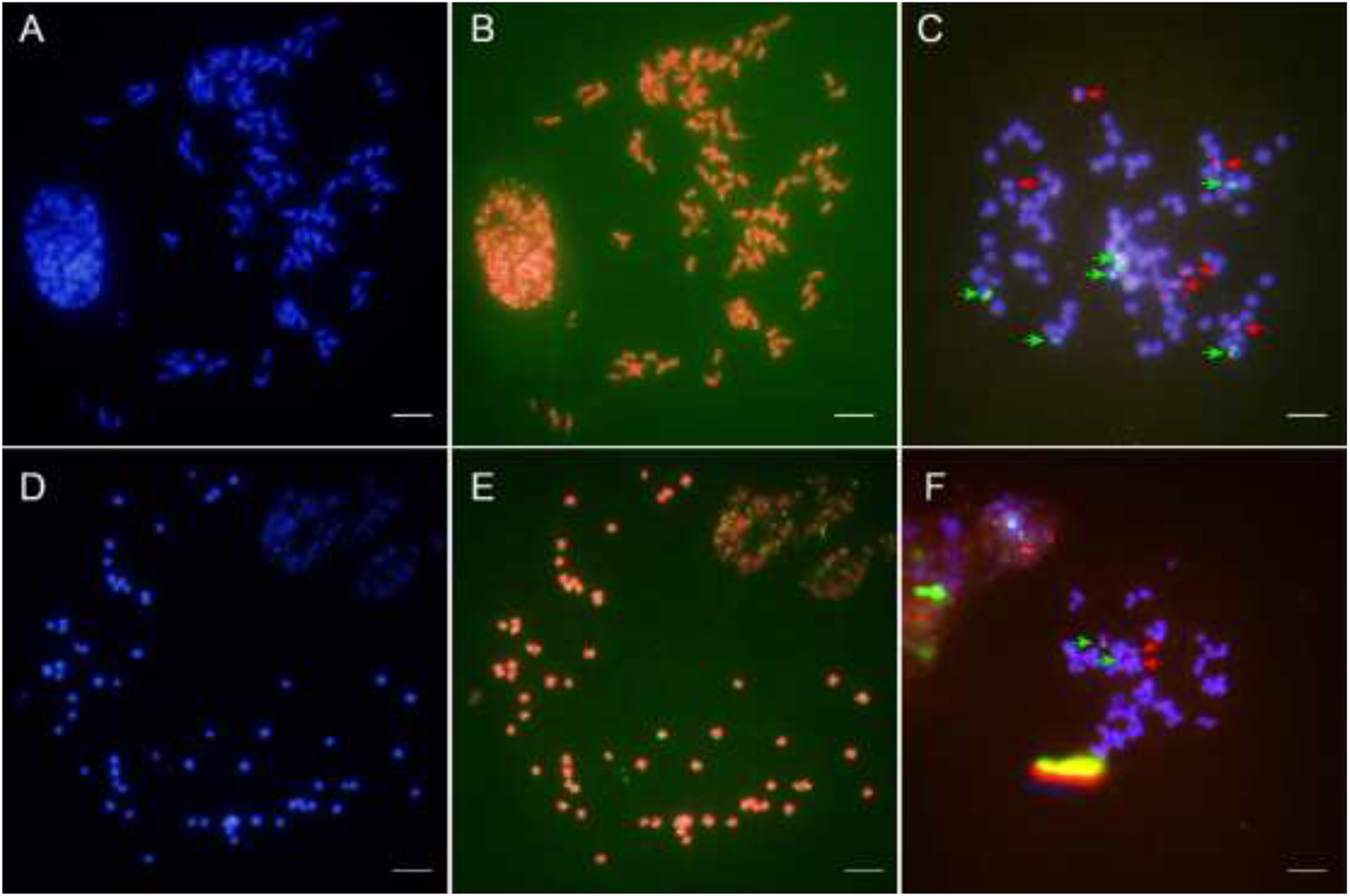
Karyotype analysis of 0004 and 0408. (A, D) DAPI staining of chromosomes of 0004 (A) and 0408 (D). (B, E) *In situ* hybridization of telomere repeat sequences of 0004 (B) and 0408 (E). (C, F) rDNA fluorescence *in situ* hybridization of chromosomes of 0004 (C) and 0408 (F). Red and green arrows denote the 5SrDNA and 18SrDNA hybridization signals. Bar = 50 μm.

### 3.2. High-quality genome assembly of A. donax

Before the genome *de novo* assembly, we conducted k-mer analysis using next generation sequencing (MGI-2000 platform). Since no whole genome sequence data of *A. donax* were available in the nucleotide sequence database, we first checked for sequence contamination by randomly selecting 100,000 reads and matched against sequences in database. About 79.8% and 84.5% reads of 0004 and 0408 were unmapped in the database, confirming the low contamination of MGI data. Besides, over 2000 reads were mapped with *Setaria viridis* and *Digitaria exilis*, indicating the homology of *A. donax* and these two species (Supplementary Figure S1A-B). The estimated genome size of 0004 and 0408 was 1.46 Gb and 3.14 Gb, respectively. In addition, we estimated the genome heterozygosity using simulated data of Arabidopsis. The heterozygosity of 0004 and 0408 genome is 0.8% and 0.3% respectively (Supplementary Figure S1C-D).

The long reads used for genome assembly were generated by PacBio platform. After quality control, we obtained 152.30 Gb reads with N50 16.92 kb for 0004 and 133.18 Gb reads with N50 21.37 kb for 0408. The preliminary assembly of 0408 was 2.97 Gb, in consistent with 3.14 Gb estimated with genome survey. However, assembly of 0004 was 2.24 Gb, much larger than 1.46 Gb. Considering the relatively high heterozygosity of 0004 genome (0.8%), we used Purge_Dups (-f .9) [28] to obtain the 1.41 Gb non-redundant genome. The GC depth analysis showed an improvement of genome quality for 0408 after redundancy reduction (Supplementary Figure S2).

We use Benchmarking Universal Single-Copy Orthologs (BUSCO) and CEGMA to assess the completeness of genome assembly. The percentage of complete BUSCOs was 99.57% and 99.69% in 0004 and 0408 genome (Supplementary Figure S3A-B), and the percentage of complete core genes was 92.74% and 91.53% in 0004 and 0408 genome (Supplementary Figure S3C-D), confirming the completeness of two genome assembly.

Besides, to determine the accuracy of the genome assembly, we mapped the MGI reads to the assemblies, the map rate is 99.60% and 99.96% for 0004 and 0408, respectively, proving the accuracy of the genome assemblies.

### 3.3. Hi-C scaffolding of A. donax genome assemblies

Since the basic chromosome basic number of *A. donax* is still unclear, we first assumed that the two *A. donax* varieties are diploid, thus the chromosome basic number is 56 and 31 for 0004 and 0408, respectively. We performed Hi-C scaffolding and the Hi-C interaction heatmap showed that both 0004 and 0408 are not diploid (Supplementary Figure S4). The number of Hi-C interaction signals for 0004 and 0408 is around 40 and 70, respectively. A hypothesis is that ancestor of all *Arundo* species is *A. plinii*, whose haploid chromosome number is 36 (n=36) [29]. Therefore, we redid Hi-C scaffolding with n = 36 for 0004 (Figure 3A). Overall, about 99.78% of total sequences were anchored into 36 pseudo chromosomes with sizes ranging from 18.58 Mb to 55.91 Mb. The final genome assembly is 1.30 Gb with Scaffold N50 37.31 Mb (Table 1 and Figure 3C). Based on these results, we speculated common *A. donax* is triploid (2n = 3x = 108), while 0004 is an aneuploid with four extra chromosomes (3x + 4 =112). The LTR Assembly Index (LAI) score of 0004 was 12.63, reaching the standard of reference quality.

**Figure 3.**
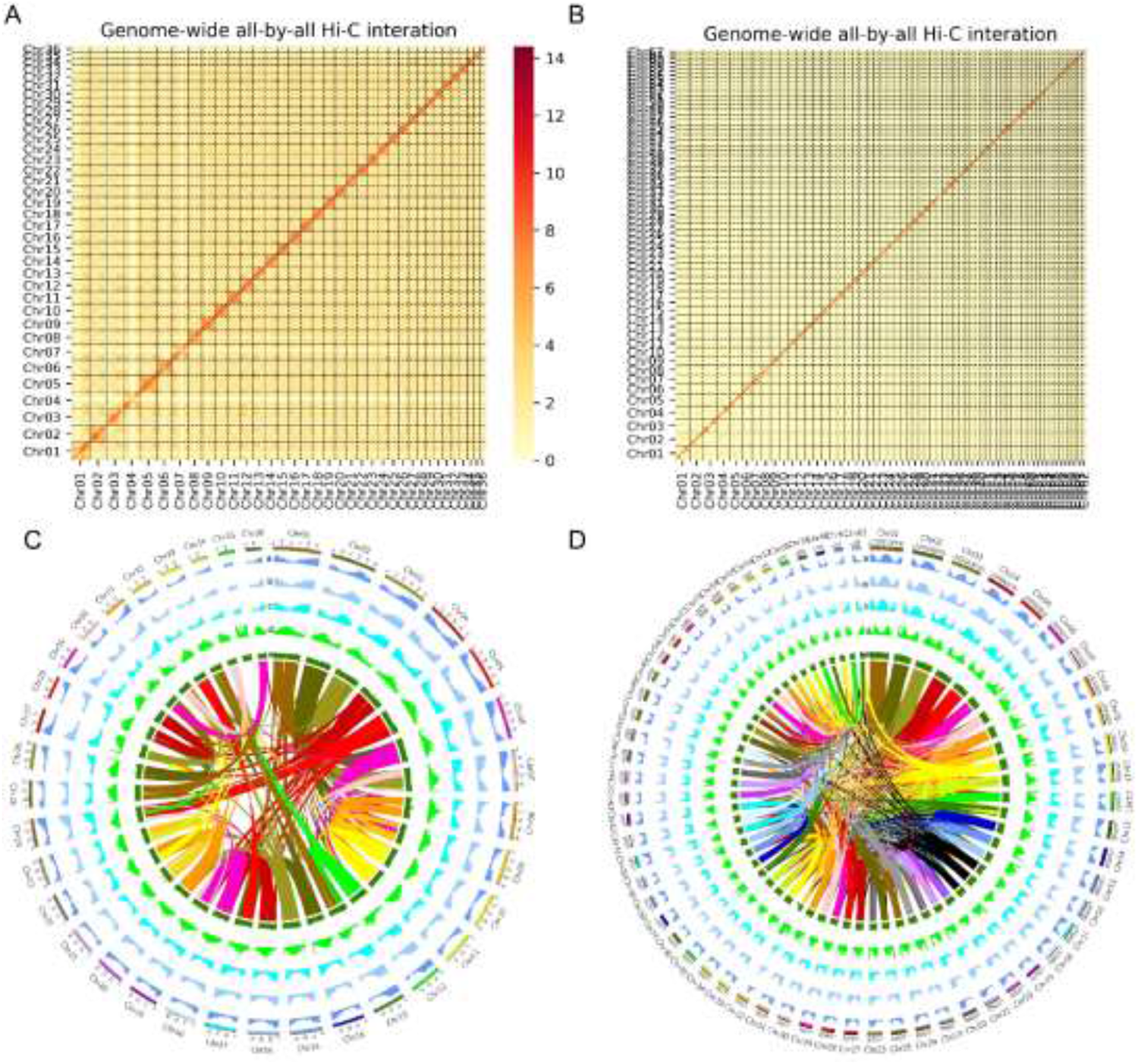
Overview of the *A. donax* genome assemblies. (A, B) Genome-wide Hi-C map of 0004 (A) and 0408 (B). (C, D) Circos plots of 0004 (C) and 0408 (D) genome assemblies. The density of GC content (a), repeated sequence distribution (b), gene density (c), functional genes (d), expressed genes (e) were calculated using 100kb non-overlap window. The innermost lines represent inter-chromosomal synteny. (E) Macrosynteny dotplot of 0004 and 0408 chromosomes.

**Table 1.** Statistics of genome assemblies.

|  | 0004 | 0408 |
| --- | --- | --- |
| **Assembly features** |  |  |
| Number of scaffolds | 65 | 162 |
| Total size of scaffolds | 1299.92 Mb | 2770.95 Mb |
| Longest scaffold | 55.91 Mb | 97.92 Mb |
| Shortest scaffold | 18.58 Mb | 24.80 Mb |
| Mean scaffold size | 20.00 Mb | 17.10 Mb |
| N50 scaffold length | 37.31 Mb | 44.86 Mb |
| L50 scaffold count | 7 | 9 |
| Scaffold GC content | 44.07% | 44.05% |
| Scaffold N content | 0.0002% | 0.0003% |
| Percentage of assembly in scaffolded contigs | 99.78% | 97.96% |
| Average number of contigs per scaffold | 1.46 | 1.12 |
| BUSCO (complete) | 99.57% | 99.69% |
| LTR Assembly Index (LAI) | 12.63 | 8.94 |
| **Gene models** |  |  |
| Number of gene models | 74,403 | 167,586 |
| Mean coding sequence length | 1192.62 | 1,138.43 |
| Mean number of exons per gene | 5.31 | 5.02 |
| Mean exon length | 224.79 | 226.57 |
| Mean intron length | 537.64 | 518.93 |
| **Non-protein-coding RNA** |  |  |
| Number of rRNA | 2,320 | 6,305 |
| Number of sRNA | 3,118 | 6,702 |
| Number of regulatory | 19 | 29 |
| Number of tRNA | 1,392 | 3,175 |

However, for the 0408 genome assembly, the chromosome basic number is obscure. The number of Hi-C interaction signals (about 70) is more than the total chromosome number (62). Therefore, we assumed that 0408 is haploid and redid Hi-C scaffolding with n = 62 for 0408 (Figure 3B). About 97.00% of total sequences were anchored into 62 pseudochromosomes with sizes ranging from 24.80 Mb to 97.92 Mb. The final genome size is 2.86 Gb with scaffold N50 44.86 Mb (Figure 3D). The LAI score of assembly constructed by Hi-C scaffolding was 8.94 for 0408, approaching the standard of reference quality.

### 3.4. Repeat elements analysis and gene model prediction

We first analyzed the interspersed repeats in 0004 and 0408 genomes. The total length of interspersed repeats is 711.24 Mb (54.71% of the genome) for 0004 and 1627.27 Mb (56.96% of the genome) for 0408. To be specific, a total of 70,102 (0.07% of the genome) and 149,490 (0.07% of the genome) simple repeat sequence (SSR) were identified in 0004 and 0408 genome; a total of 89,378 (0.71% of the genome) and 181,468 (0.78% of the genome) tandem repeat sequences were identified in 0004 and 0408 genome; a total of 1,400,366 (51.54% of the genome) and 2,937,325 (52.91% of the genome) transposable elements (TEs) were identified in 0004 and 0408 genome. The detailed statistics of TEs were listed in Supplementary file2 (TE).

We performed gene structure prediction by combining transcriptome prediction, homologous protein prediction, and ab initio prediction. Firstly, we found that the mapped rate of RNA-seq data to the genome in four tissues were all over 90% (Supplementary Table 1), and the mapped rate of Pacbio Isoseq to the genome is 99.86% for 0004 and 99.87% for 0408, further confirming the accuracy of transcriptome data and the genome assemblies. The RNA-seq and Pacbio transcripts were used for gene prediction, resulting in 49,524 / 80,249 predicted genes for 0004 and 0408 respectively. Secondly, we selected five Poaceae plants, including *Saccharum spontaneum*, *Sorghum bicolor*, *Zea mays*, *Triticum aestivum* and *Oryza sativa* for homologous protein prediction. A total of 94,613 / 172,436 genes for 0004 and 0408 were predicted. Thirdly, we performed ab initio prediction, and 78,102 / 174,926 gene models for 0004 and 0408 were predicted. The final gene set was obtained by integrating the above results. In total, 74,403 / 167,586 gene models with average gene length 3,507.41 / 3,226.92 bp, average CDS length 1192.62 / 1,138.43 bp, average exon length 224.79 / 226.57 bp, average intron length 537.64 / 518.93 bp for 0004 and 0408 respectively (Supplementary file2 (Gene prediction)). Except the gene model, we also predicted non-coding RNA. In total, 2,320 / 6,305 rRNA, 3,118 / 6702 small RNA, 1,392 / 3,175 tRNA were identified in 0004 and 0408 genome. The parameters of the two assemblies were listed in Supplementary file2 (ncRNA).

### 3.5. Gene function annotation and evaluation of genome annotation

We predicted the gene function based on five databases, including Non-Reduntant Protein Database (NR), Kyoto Encyclopedia of Gene and Genomes (KEGG), Eukaryotic Orthologous Groups of protein (KOG), GO and Swissprot (Supplementary Figure S5). In total, 67,377 / 152,275 genes were annotated, accounting for 90.56% / 90.86 % of the 0004 and 0408 genomes (Supplementary file2 (Gene prediction)). The co-annotated gene number is 16,877 / 37,776 for 0004 and 0408 genomes (Supplementary Figure S6).

The annotated gene sets were evaluated using BUSCO. Among the 1,614 BUSCO groups, about 98.45% / 97.96% of complete gene elements can be found in the annotated gene set for 0004 and 0408 respectively, indicating that the majority of conservative gene predictions are relatively complete, indicating the high reliability of the gene prediction result. Besides, the proportion of expressed genes in four tissues ranged from 71.09% to 80.34% in 0004 and 67.01% to 73.75% in 0408. Total expressed genes account for 86.60% and 82.53% of the whole gene sets of 0004 and 0408, respectively (Supplementary file2 (Transcripts)). The gene structure in 0004 and 0408 genome showed similar distribution trend with other Poaceae plants, including gene length, CDS length, exon length, exon number, intron length and intron number (Supplementary Figure S7), demonstrating the reliability of the genome annotations.

### 3.6. Comparative genomic analysis

To clarify the relationship between 0004 and 0408, we compared the genomes of 0004 and 0408. We identified some chromosome structure variations of the two genomes. More interestingly, the two genomes showed an obvious 1:2 pattern of chromosome alignment between 0004 and 0408 (Supplementary Figure S8), hinting that two subgenomes exist in the 0408genome. We used SubPhaser [65] to differentiate the two subgenomes of 0408, heatmap showed that subgenome A has 28 chromosomes and subgenome B has 34 chromosomes (Figure 4B). Collinearity analysis using gene CDS sequence revealed 3:6 ratio of 0004:0408 (Supplementary Figure S9) and 3:3:3 ratio of 0004:0408sbA:0408sbB (Figure 4A) in chromosome segments.

**Figure 4.**
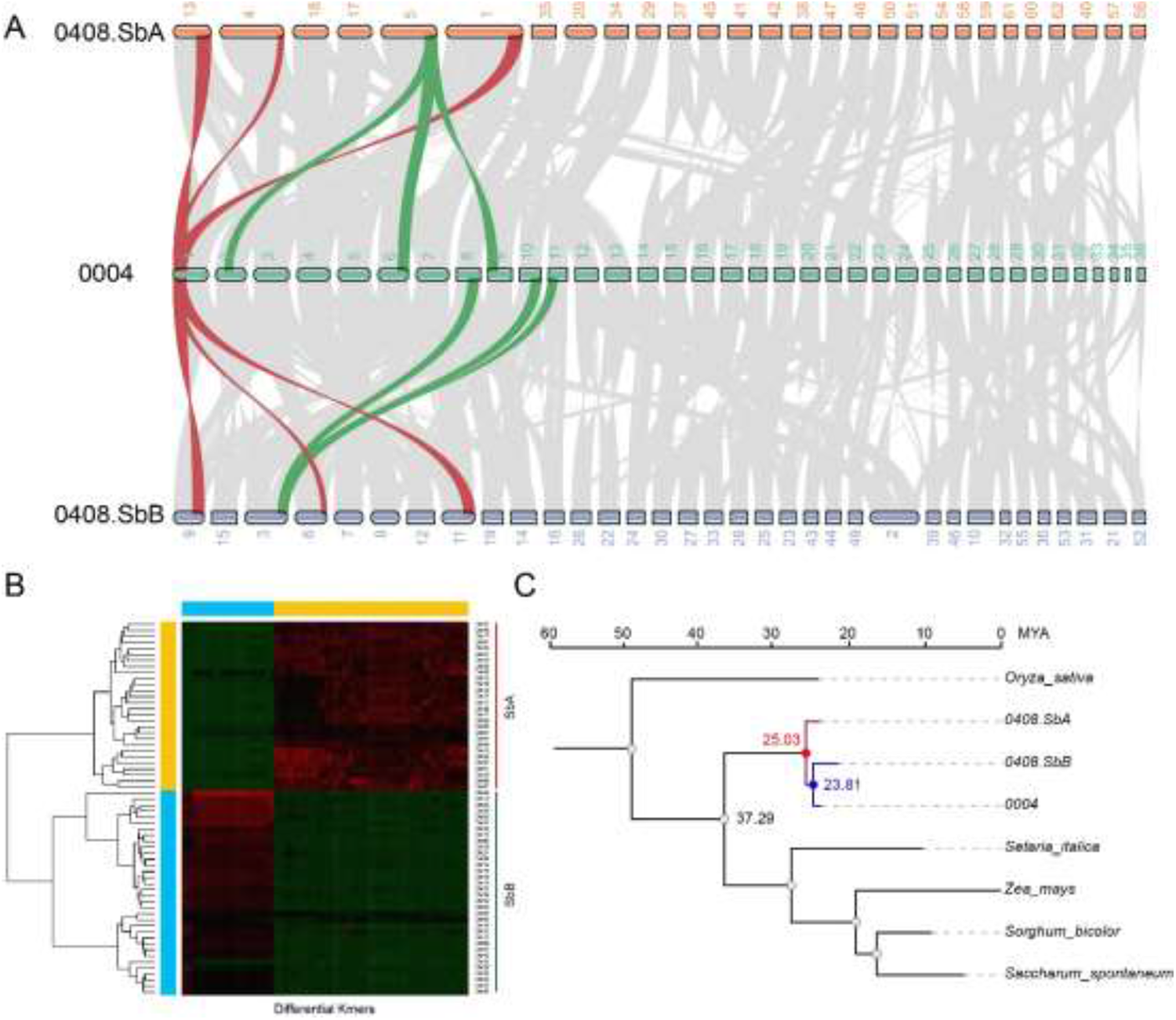
Comparative genomics analysis of 0004 and 0408. (A) Syntenic relationships between 0004 and subgenomes of 0408. Red and green stripes represent the correspondence of chromosome segments between 0004 and subgenomes of 0408. (B) Clustering based on specific K-mers to partition the two subgenomes of 0408. Red and green chromosomes correspond to the A and B subgenome (C) Timetree of *Arundo* and several other Gramineae plants. The evolutionary relationships between species were estimated by highly conserved single-copy genes. MYA, million years ago.

To investigate the genome evolutionary history of *A. donax*, gene family clustering was carried out using 0004/0408 and five other Gramineae species. From the gene family clustering, single-copy orthologs shared by *Arundo* and five other plants were used for phylogenetic analysis and divergence time estimation, which showed that subfamily *Arundo* and *Paniceae* shared a common ancestor ∼37.29 Ma. The divergence time of 0004 and 0408 was 25.03 MYA. Besides, subgenome B showed higher similarity with 0004 than that of subgenome A (Figure 4C).

Consider the relatively close relationship between *S. italica* and *A. donax*, we carried out further collinearity analysis and revealed 3:1 ratio of *S. italica*:0004 (Supplementary Figure S10A), and 1:6 ratio of *S. italica*:0408 (Supplementary Figure S10B) in chromosome segments. Based on these results, we speculated common *A. donax* is an alloenneaploid (3n = 9x = 108) and emerald *A. donax* is an allohexaploid (2n = 6x = 72). We use SubPhaser to further split the subgenomes of 0004, the heatmap showed that the chromosomes were clustered to two groups, in which subgenome A has 12 chromosomes and Subgenome B has 24 chromosomes (Supplementary Figure S11).

### 3.7 Gene family analysis of A. donax

A total of 36,390/31,024 gene clusters were identified in 0004/0408 genomes, in which 10,468 clusters were unique to *A. donax*, and 297/4425 gene clusters were unique to 0004/0408, respectively (Figure 5A). *A. donax* undergone dramatic gene family expansion during evolution (Figure 5B). The 3835 expanded gene families were enriched in GO terms like “response to water deprivation”, “response to oxidative stress”, “response to osmotic stress”, “response to cold” and “response to heat” (Figure 5C), hinting that *A. donax* emergenced from the grass family because of the extreme environment in earth.

**Figure 5.**
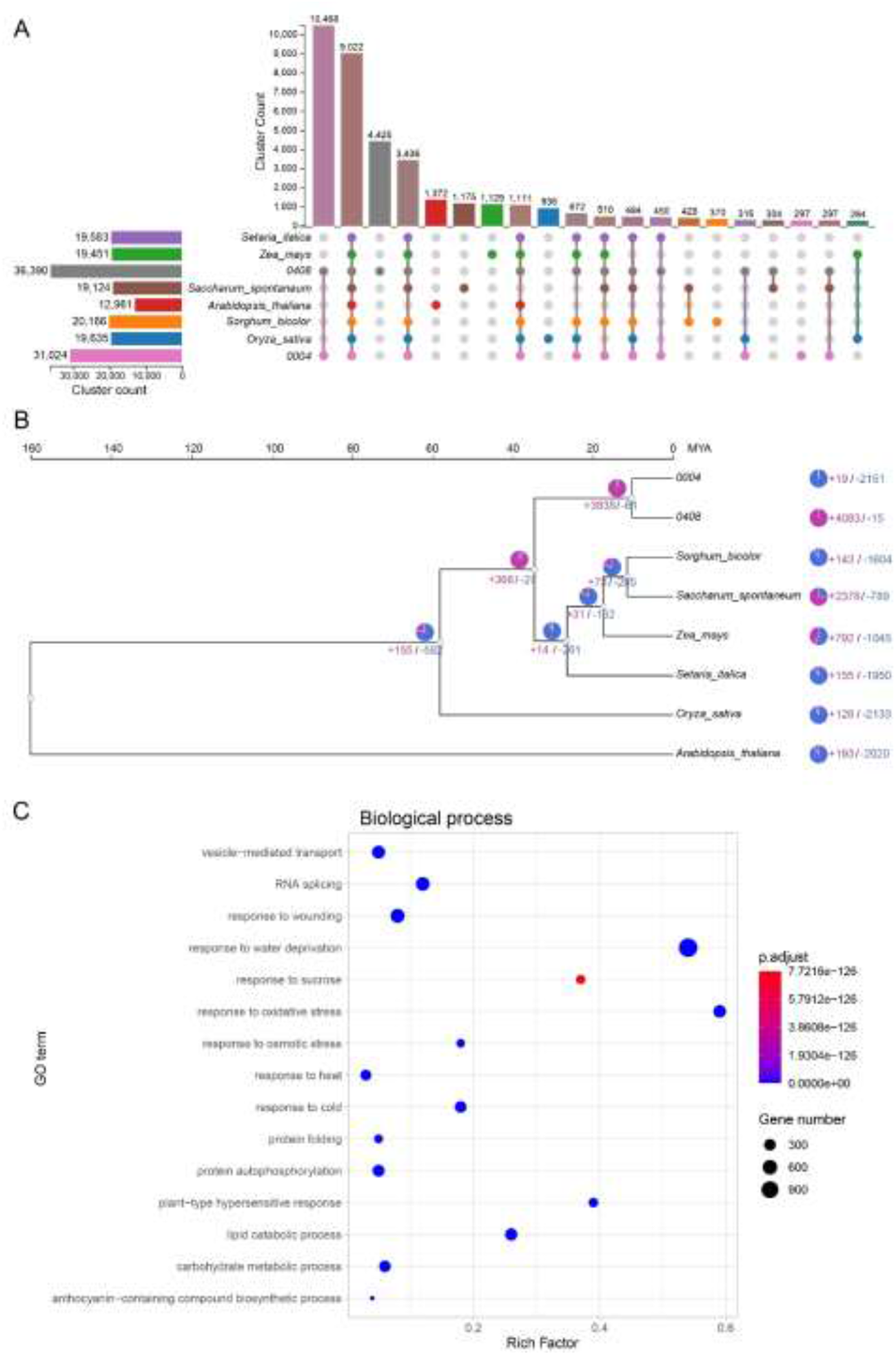
Gene family contraction and expansion analysis of 0004 and 0408. (A) The UpSet table displays the count of orthologous clusters for each species, along with the count of unique gene clusters and the count of shared gene clusters among different species. (B) A pie chart was utilized to visualize gene families with altered gene numbers, representing the contracted gene families (depicted in purple) and expanded gene families (depicted in blue). (C) GO enrichment of 3835 expanded gene clusters of *A. donax* (top 15 of biological process category). The color of dot denotes the adjusted p-value using hypergeometric test. The size of dot denotes the gene number of GO terms.

We also analyze the whole-genome duplication (WGD) events in both 0004 and 0408 genomes (Supplementary Figure S12). For 0004, except for the ancient ρ event in Poaceae, a recent burst WGD event occurred ∼13.5 MYA. While for 0408, the ancient WGD is hardly detectable in the whole genome, and two recent burst WGD events occurred about ∼13.1 MYA and ∼5.4 MYA. We speculate that the WGD events occurred ∼13 MYA is the common WGD event in *A. donax*, and emerald *A. donax* undergone a unique WGD event.

## 4. Discussion

### 4.1 A. donax is a promising energy crop with complex genome

The rapidly deteriorating environment on earth has made an urgent claim for carbon emission reduction [64]. Using bioenergy to replace part of fossil energy can make remarkable contributions to reducing carbon emissions [65]. However, land-intensive bioenergy is limited by the finite land resource. Especially in China, because of the huge population size, croplands are inviolable to guarantee food security. However, there are about 78 million hectares of marginal lands except for the cropland [66]. Due to climate, terrain, soil and other limiting factors, marginal lands cannot be planted with food crops, while energy crops can often grow on these lands owing to their great environmental adaptability. Of all the energy crops, the high yield, extensive adaptability and low inputs make *A. donax* an ideal energy crop.

Here, we generated the first two chromosome-scale assemblies of the *A. donax* genome using HiFi reads and Hi-C technology. According to the LAI score (12.63) of the assemblies, the assembly of 0004 reached the criterion of reference quality (Table 1). Based on the genomic synteny, we speculated that common *A. donax* is an alloenneaploid (3n = 9x = 108), and the chromosome component is AAABBBBBB. The aneuploid 0004 was generated because of the abnormal replication of four chromosomes (3n + 4 = 112). In contrast, emerald *A. donax* is an allohexaploid (2n = 6x = 72) and the chromosome component is AAABBB. The aneuploid 0408 was generated because of the abnormal deletion of ten chromosomes (2n – 10 = 62). Overall, the genome assembly of 0408 did not reach to the reference quality (LAI score = 8.63), and the Hi-C heatmap also showed that some chromosomes can be further split into two segments like chromosome 1, 2, 3, 4, 5, 10 etc (Figure 3B). Therefore, to obtain a more complete genome assembly of emerald *A. donax*, an euploid (2n = 72) plant is needed to reassembly the genome.

### 4.2 The reference genome will contribute to genetic improvement of A. donax

Clonal propagation crops like potatoes often showed small genetic gains. Using precise genome design, vigorous F_1_ hybrids potatoes has been developed recently [67]. Similar to potato, *A. donax* is completely agamic and shows a narrow genetic variability. To accelerate the genetic improvement of *A. donax* in the future, we need to elucidate the mechanism of infertility of *A. donax*. In addition, the genetic transformation of *A. donax* encounters difficulties, and no transgenic *A. donax* has been truly reported until now. Several developmental regulators, including BBM, WUS, GRF4 and GIF were reported to improve transformation efficiency in crops like corn and wheat [68]. Therefore, cloning key developmental regulators in *A. donax* may help to break down technical barriers in genetic transformation of *A. donax*. To overcome the above two difficulties, a reference genome is indispensable.

Until now, little is known about the *A. donax* genome, except for a few transcriptome studies [69–74]. Our result showed that *A. donax* has a complex genome that contains a large number of genes, many of which are involved in important biological processes such as growth, development, and stress response. Mining important genes from the reference genome will undoubtedly contribute to the genetic improvement of *A. donax*. Future research on *A. donax* will continues to uncover new insights into the plant’s biological properties and the potential for its utilization in different industries. It will undoubtedly uncover more valuable information and offer even more exciting opportunities to take advantage of this remarkable plant species.

## Author Contributions

Daohong Wu and Hai Peng: Project design, writing review and editing. Mengmeng Ren: Data analysis and writing. Fupeng Liu and Xiaohong Han: Plant material collection. All authors have read and agreed to the published version of the manuscript.

## Institutional Review Board Statement

Not applicable.

## Informed Consent Statement

Not applicable.

## Data Availability Statement

The data presented in this study are available on request from the corresponding author. Raw sequence data will be available on line later.

## Acknowledgments

Not applicable.

## References

1. Corno, L.; Pilu, R.; Adani, F. Arundo donax L.: a non-food crop for bioenergy and bio-compound production. Biotechnol. Adv. 2014, 32, 1535–1549.

2. Jámbor A.; Török, A. The Economics of *Arundo donax*—A Systematic Literature Review. Sustainability 2019, 11, 4225.

3. Pilu, R.; Manca, A.; Landoni, M. *Arundo donax* as an energy crop: pros and cons of the utilization of this perennial plant. Maydica 2013, 58.

4. Zhang, J.; Li, Y.; Zhang, C.; Jing, Y. Adsorption of malachite green from aqueous solution onto carbon prepared from *Arundo donax* root. J. Hazard. Mater. 2008, 150, 774–782.

5. Angelini, L.G., Ceccarini, L.; Bonari, E. Biomass yield and energy balance of giant reed (*Arundo donax* L.) cropped in central Italy as related to different management practices. Eur. J. Agron. 2005, 22, 375–389.

6. Angelini, L.G.; Ceccarini, L.; Nasso, N.; Bonari, E. Comparison of *Arundo donax* L. and *Miscanthus* x *giganteus* in a long-term field experiment in Central Italy: Analysis of productive characteristics and energy balance. Biomass & Bioenergy 2009, 33, 635–643.

7. Nasso, N.N.O.D.; Roncucci, N.; Bonari, E. Seasonal Dynamics of Aboveground and Belowground Biomass and Nutrient Accumulation and Remobilization in Giant Reed (*Arundo donax* L.): A Three-Year Study on Marginal Land. Bioenergy Research 2013, 6, 725–736.

8. Nackley, L.L.; Kim, S.H. A salt on the bioenergy and biological invasions debate: salinity tolerance of the invasive biomass feedstock *Arundo donax*. GCB Bioenergy 2015, 7, 752–762.

9. Sánchez, E.; Scordia, D.; Lino, G. Salinity and Water Stress Effects on Biomass Production in Different *Arundo donax* L. Clones. BioEnergy Res. 2015, 8, 1461–1479.

10. Mirza, N.; Mahmood, Q.; Pervez, A.; Ahmad, R.; Farooq, R.; Shah, M.M.; Azim, M.R. Phytoremediation potential of *Arundo donax* in arsenic-contaminated synthetic wastewater. Bioresource Technol. 2010, 101, 5815–5819.

11. Mirza, N.; Pervez, A.; Mahmood, Q.; Shah, M.M.; Shafqat, M.N. Ecological restoration of arsenic contaminated soil by *Arundo donax* L. Ecol. Eng. 2011, 37, 1949–1956.

12. Papazoglou, E.G.; Karantounias, G.A.; Vemmos, S.N.; Bouranis D.L. Photosynthesis and growth responses of giant reed (*Arundo donax* L.) to the heavy metals Cd and Ni. Environ. Int. 2005, 31, 243–249.

13. Ahmad, R.; Liow, P.S.; Spencer, D.F.; Jasieniuk, M. Molecular evidence for a single genetic clone of invasive *Arundo donax* in the United States. Aquat. Bot. 2008, 88, 113–120.

14. Danelli, T.; Laura, M.; Savona, M.; Landon, M.; Adani, F.; Pilu, R. Genetic Improvement of *Arundo donax* L.: Opportunities and Challenges. Plants 2020, 9, 1584.

15. Malone, J.M.; Virtue, J.G.; Williams, C.; Preston, C. Genetic diversity of giant reed (*Arundo donax*) in Australia. Weed Biol. Manag. 2017, 17.

16. Tarin, D.; Pepper, A.E.; Goolsby, J.A.; Moran, P.J.; Arquieta, A.C.; Kirk, A.E.; Manhart, J.R. Microsatellites Uncover Multiple Introductions of Clonal Giant Reed (*Arundo donax*). Invas. Plant Sci. Mana. 2013, 6, 328–338.

17. Bucci, A.; Cassani, E.; Landoni, M.; Cantaluppi, E.; Pilu, R. Analysis of chromosome number and speculations on the origin of *Arundo donax* L. (Giant Reed). Cytolo. Genet. 2013, 47, 237–241.

18. Mariani, C.; Cabrini, R.; Danin, A.; Piffanelli, P.; Fricano, A.; Gomarasca, S.; Dicandilo, M.; Grassi, F.; Soave, S. Origin, diffusion and reproduction of the giant reed (*Arundo donax* L.): a promising weedy energy crop. Ann. Appl. Biol. 2010, 157, 191–202.

19. Clevering, O.A.; Lissner, J. Taxonomy, chromosome numbers, clonal diversity and population dynamics of *Phragmites australis*. Aquat. Bot. 1999, 66, 249–250.

20. Hunter, A.W.S. A Karyosystematic investigation in Gramineae. Can. J. Res. 1934, 11, 213–241.

21. Pizzolongo, P. Osservazioni cariologiche su *Arundo donax* e *Arundo plinii*. Annuali Bot. 1962, 27, 173–187.

22. Christopher, J.; Abraham, A. Studies on the cytology and phylogeny of South Indian grasses I. Subfamilies Bambusoideae, Oryzoideae, Arundinoideae and Festucoideae. Cytologia 1971, 36, 579– 594.

23. Haddadchi, A.; Gross, C.L.; Fatemi, M. The expansion of sterile *Arundo donax* (Poaceae) in southeastern Australia is accompanied by genotypic variation. Aquat. Bot. 2013, 104, 153–161.

24. Wang, Y.; Yu, J.; Jiang, M.; Lei, W.; Zhang, X.; Tang, H. Sequencing and Assembly of Polyploid Genomes. Methods Mol. Biol. 2023, 2545, 429–458.

25. Peng, Y., Yan, H., Guo, L., Deng, C.; Wang, C., Wang, Y., Kan, L.; Zhou, P.; Y, K.; Dong, X.; Liu, X.; Su, Z.; Peng, Y.; Zhao, J.; Deng, D.; Xu, Y.; Li, Y.; Jiang, Q.; Li, Y.; Wei, L.; Wang, J.; Ma, J.; Hao, M.; Li, W.; Kang, H.; Peng, Z.; Liu, D.; Jia, J.; Zheng, Y.; Ma, T.; Wei, Y.; Lu, F.; Ren, C. Reference genome assemblies reveal the origin and evolution of allohexaploid oat. Nat. Genet. 2022, 54, 1248– 1258.

26. Sun, H.; Jiao, W.B.; Krause, K.; Campoy, J.A.; Goel, M.; Folz-Donahu, K.; Kukat, C.; Huettel, B.; Schneeberger, K. Chromosome-scale and haplotype-resolved genome assembly of a tetraploid potato cultivar. Nat. Genet. 2022, 54, 342–348.

27. Chen, H.; Zeng, Y.; Yang, Y.; Huang, L.; Tang, B.; Zhang, H.; Hao, F.; Liu, W.; Li, Y.; Liu, Y.; Zhang, X.; Zhang, R.; Zhang, Y.; Li, Y.; Wang, K.; He, H.; Wang, Z.; Fan, G.; Yang, H.; Bao, A.; Shang, Z.; Chen, J.; Wang, W.; Qiu Q. Allele-aware chromosome-level genome assembly and efficient transgene-free genome editing for the autotetraploid cultivated alfalfa. Nat. Commun. 2020, 19, 2494.

28. Bayani, J.; Squire, J.A. Fluorescence *in situ* Hybridization (FISH). Current Protocols in Cell Biology 2004, Chapter 22.

29. Jiang, J. Fluorescence *in situ* hybridization in plants: recent developments and future applications. Chromosome Res. 2019, 27, 153–165.

30. Dolezel, J.; Greilhuber, J.; Suda, J. Estimation of nuclear DNA content in plants using flow cytometry. Nat. Protoc. 2007, 2, 2233–2244.

31. Kokot, M.; Dlugosz, M.; Deorowicz, S. KMC 3: counting and manipulating k-mer statistics. Bioinformatics 2017, 33, 2759–2761.

32. Sun, H.; Ding, J.; Piednoël, M.; Schneeberge, K. FindGSE: Estimating genome size variation within human and Arabidopsis using k -mer frequencies. Bioinformatics 2018, 34: 550–557.

33. Cheng, H., Concepcion, G.T.; Feng X.; Zhang, H.; Li, H. Haplotype-resolved *de novo* assembly using phased assembly graphs with hifiasm. Nat. Methods 2021, 18, 1–6.

34. Simão, F.A.; Waterhouse, R.M.; Ioannidis, P.; Kriventseva, E.V.; Zdobnov, E.M. BUSCO: assessing genome assembly and annotation completeness with single-copy orthologs. Bioinformatics 2015, 31, 3210–3212.

35. Parra, G.; Bradnam, K.; Korf, I. CEGMA: a pipeline to accurately annotate core genes in eukaryotic genomes. Bioinformatics 2007, 23, 1061–1067.

36. Li, H.; Durbin, R. Fast and accurate long-read alignment with Burrows-Wheeler transform. Bioinformatics 2010, 26, 589–595.

37. Li, H.; Handsaker, B. Wysoker A, et al. The Sequence Alignment/Map format and SAMtools. Bioinformatics 2009, 25: 2078–2079.

38. Danecek, P.; McCarthy, S.A. BCFtools/csq: haplotype-aware variant consequences. Bioinformatics 2017, 33, 2037–2039.

39. Ou, S.; Chen, J.; Ning, J. Assessing genome assembly quality using the LTR Assembly Index (LAI). Nucleic Acids Res. 2018, 46, e126.

40. Servant, N.; Varoquaux, N.; Lajoie, B.R.; Viara, E.; Chen, C.; Vert, J.P.; Heard, E.; Dekker, J.; Barillot, E. HiC-Pro: an optimized and flexible pipeline for Hi-C data processing. Genome Biol. 2015, 16, 259.

41. Chen, S.; Zhou, Y.; Chen, Y.; Gu, J. fastp: an ultra-fast all-in-one FASTQ preprocessor. Bioinformatics 2018, 34, i884–i890.

42. Langmead, B.; Salzberg, S. Fast gapped-read alignment with Bowtie 2. Nat. Methods 2012, 9, 357–359.

43. Burton, J.N.; Adey, A.; Patwardhan, R.P.; Qiu, R.; Kitzman, J.O.; Shendure, J. Chromosome-scale scaffolding of *de novo* genome assemblies based on chromatin interactions. Nat. Biotechnol. 2013, 31, 1119–1125.

44. Wang, X.; Wang, L. GMATA: An Integrated Software Package for Genome-Scale SSR Mining, Marker Development and Viewing. Front. Plant Sci. 2016, 7, 1350.

45. Benson, G. Tandem repeats finder: a program to analyze DNA sequences. Nucleic Acids Res. 1999, 27, 573–580.

46. Han, Y.; Wessler, S.R. MITE-Hunter: a program for discovering miniature inverted-repeat transposable elements from genomic sequences. Nucleic Acids Res. 2010, 38: e199.

47. Xu, Z.; Wang, H. LTR_FINDER: an efficient tool for the prediction of full-length LTR retrotransposons. Nucleic Acids Res. 2007, 35 (Web Server issue), W265–W268.

48. Ellinghaus, D.; Kurtz, S.; Willhoeft, U. LTRharvest, an efficient and flexible software for *de novo* detection of LTR retrotransposons. BMC Bioinformatics 2008, 9, 18.

49. Ou, S.; Jiang, N. LTR_retriever: A Highly Accurate and Sensitive Program for Identification of Long Terminal Repeat Retrotransposons. Plant Physiol. 2018, 176, 1410–1422.

50. Chen, N.S. Using RepeatMasker to identify repetitive elements in genomic sequences. Curr. Protoc. Bioinformatics 2004, 4, Unit 4.10.

51. Keilwagen, J.; Wenk, M.; Erickson, J.L.; Schattat, M.H.; Grau, J.; Hartung, F. Using intron position conservation for homology-based gene prediction. Nucleic Acids Res. 2016, 44, e89.

52. Dobin, A.; Davis, C.A.; Schlesinger, F., Jorg, Drenkow.; Zaleski, C.; Jha, S.; Batut, P.; Chaisson, M.; Gingeras, T.R. STAR: ultrafast universal RNA-seq aligner. Bioinformatics 2013, 29, 15–21.

53. Kovaka, S.; Zimin, A.V.; Pertea, G.M.; Razaghi, R.; Salzberg, S.L.; Pertea, M. Transcriptome assembly from long-read RNA-seq alignments with StringTie2. Genome Biol. 2019, 20, 278.

54. Haas, B.J.; Salzberg, S.L.; Zhu, W.; Pertea, W.; Allen, J.E.; Orvis, J.; White, O.; Buell, C.R.; Wortman, J.R. Automated eukaryotic gene structure annotation using EVidenceModeler and the Program to Assemble Spliced Alignments. Genome Biol. 2008, 9, R7.

55. Mario, S.; Burkhard, M. AUGUSTUS: a web server for gene prediction in eukaryotes that allows user-defined constraints. Nucleic Acids Res. 2005, 33 (Web Server issue), W465–W467.

56. Haas, B.J.; Salzberg, S.L.; Zhu, W.; Pertea, M.; Allen, J.E; Orvis, J.; White, O.; Buell, C.R.; Wortman, J.R. Automated eukaryotic gene structure annotation using EVidenceModeler and the Program to Assemble Spliced Alignments. Genome. Biol. 2008, *R7*.

57. Zdobnov, E.M.; Apweiler, R. InterProScan--an integration platform for the signature-recognition methods in InterPro. Bioinformatics 2001, 17, 847–848.

58. Chan, P.P.; Lin, B.Y.; Mak, A.J. tRNAscan-SE: a program for improved detection of transfer RNA genes in genomic sequence. Nucleic Acids Res. 1997, 25, 955–964.

59. Nawrocki, E.P.; Eddy, S.R. Infernal 1.1: 100-fold faster RNA homology searches. Bioinformatics 2013, 29, 2933–2935.

60. Lagesen, K.; Hallin, P.; Rødland, E.A.; Stærfeldt, H.H.; Rognes, T.; Ussery, D.W. RNAmmer: consistent and rapid annotation of ribosomal RNA genes. Nucleic Acids Res. 2007, 35, 3100–3108.

61. Li, H. Minimap2: pairwise alignment for nucleotide sequences. Bioinformatics 2018, 34, 3094–3100.

62. Sun J, Lu F, Luo Y, Bie L, Xu L, Wang Y. OrthoVenn3: an integrated platform for exploring and visualizing orthologous data across genomes. Nucleic Acids Res. 2022, 51(*W1*), W397–W403.

63. Sun P, Jiao B, Yang Y, Shan L, Li T, Li X, Xi Z, Wang X, Liu J. WGDI: A user-friendly toolkit for evolutionary analyses of whole-genome duplications and ancestral karyotypes. Mol. Plant 2022, 15, 1841–1851.

64. Guan, D.; McCarthy, S.A.; Wood, J.; Howe, K.; Wang, Y.; Durbin, R. Identifying and removing haplotypic duplication in primary genome assemblies. Bioinformatics 2020, 36, 2896–2898.

65. Patz, J.A.; Frumkin, H.; Holloway, T.; Vimont, D.J.; Haines, A. Climate Change:challenges and opportunities for global health. JAMA 2014, 312, 1565–1580.

66. Walter, V.R.; Mariam, K.A.; Christopher B.F. The future of bioenergy. Glob Chang Biol. 2020, 26, 274–286.

67. Jia, K., Wang, Z., Wang, L., Li, G., Zhang, W., Wang, X., Xu, F., Jiao, S., Zhou, S., Liu, H., Ma, Y., Bi, G., Zhao, W., El-Kassaby, Y. A., Porth, I., Li, G., Zhang, R., Mao, J. SubPhaser: a robust allopolyploid subgenome phasing method based on subgenome-specific k-mers. New Phytol. 2022, 235, 801–809.

68. Tang, Y.; Xie, J.S.; Geng, S. Marginal Land-based Biomass Energy Production in China. J. Integr. Plant Biol. 2010, 52, 112–21.

69. Zhang, C.; Yang, Z.; Tang, D.; Zhu, Y.; Wang, P.; Li, D.; Zhu, G.; Xiong, X.; Shang, Y.; Li, C.; Huang, S. Genome design of hybrid potato. Cell 2021, 184, 3873–3883.e12.

70. Chen, Z.; Debernardi, J.M.; Dubcovsky, J.; Gallavotti, A. Recent advances in crop transformation technologies. Nat. Plants 2022, 8, 1343–1351

71. Evangelistella, C.; Valentini, A.; Ludovisi, R.; Firrincieli, A.; Fabbrini, F.; Scalabrin, S.; Cattonaro, F.; Morgante, M.; Mugnozza, G.S.; Keurentjes, J.J.B.; Harfouche, A. *De novo* assembly, functional annotation, and analysis of the giant reed (*Arundo donax* L.) leaf transcriptome provide tools for the development of a biofuel feedstock. Biotechnol. Biofuels 2017, 10, 138.

72. Fu, Y.; Poli, M.; Sablok, G.; Wang, B.; Liang, Y.; Porta, N.L.; Velikova, V.; Loreto, F., Li, M., Varotto, C. Dissection of early transcriptional responses to water stress in *Arundo donax* L. by unigene-based RNA-seq. Biotechnol. Biofuels 2016, 9, 54.

73. Sablok, G.; Fu, Y.; Bobbio, V.; Laura, M.; Rotino, G.L.; Bagnaresi, P.; Allavena, A.; Velikova, V.; Viola, R.; Loreto, F.; Li, M.; Varotto, C. Fuelling genetic and metabolic exploration of C3 bioenergy crops through the first reference transcriptome of *Arundo donax* L. Plant Biotechnol. J. 2014, 12, 554–567.

74. Sicilia, A.; Testa, G.; Santoro, D.F.; Cosentino, S.L.; Piero, A.R.L. RNASeq analysis of giant cane reveals the leaf transcriptome dynamics under long-term salt stress. BMC Plant Biol. 2019, 19.

