## Supplementary material for "Chromosome-scale Genome Assembly of the Allopolyploid *Arundo donax*": ~WRL0731.tmp

**Supplementary Figures**

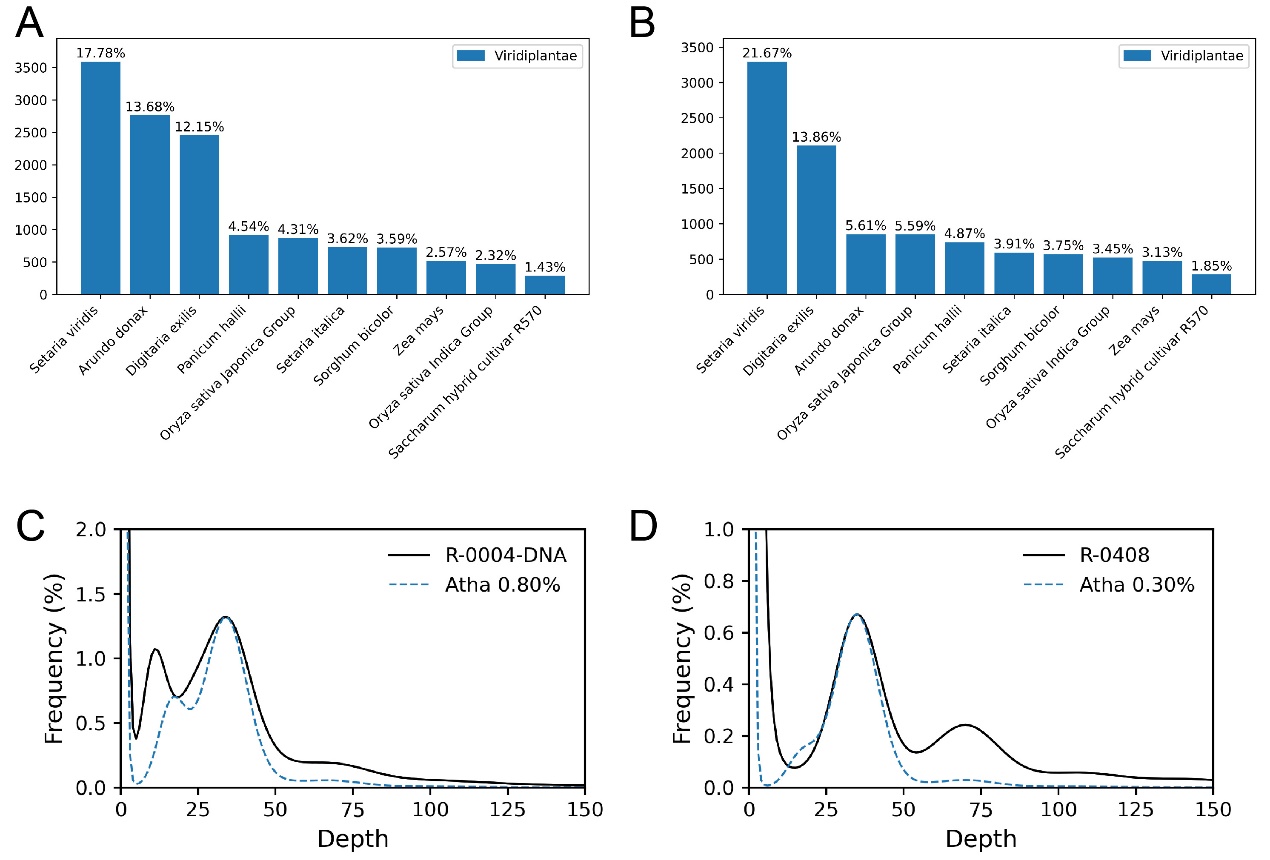

**Supplementary Figure S1.** Genome survey of 0004 and 0408. (A-B) Main species mapped in nucleotide sequence database using MGI sequence of 0004 (A) and 0408 (B). (C-D) K-mer distribution curve and heterozygosity simulation curve of 0004 (C) and 0408 (D). Horizontal axis is the K-mer depth, vertical axis is frequency of the K-mer depth. Genome of *Arabidopsis thaliana* (Atha) was used to estimate the genome heterozygosity of *A.donax*.

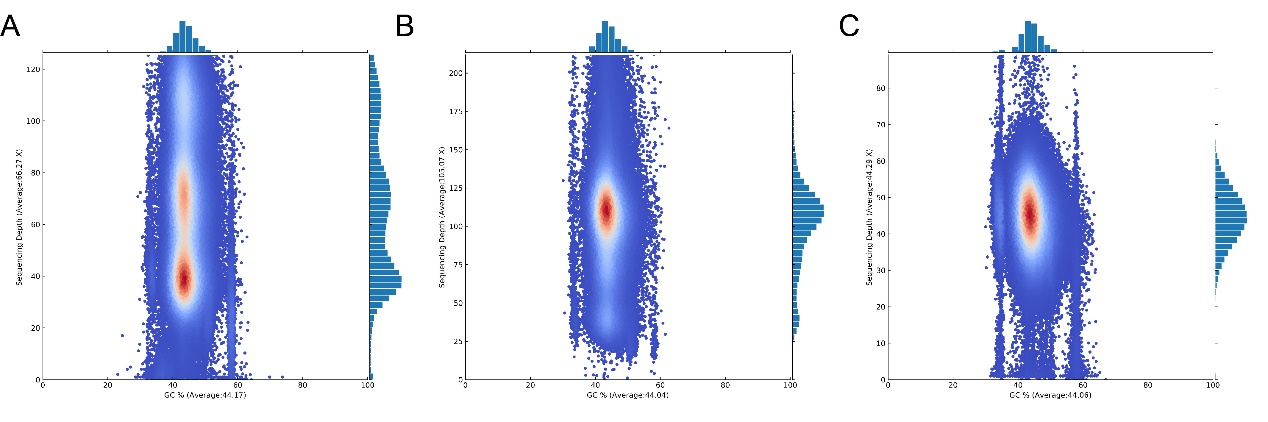

**Supplementary Figure S2.** GC depth distribution of 0004 genome before elimination of redundancy (A), 0004 genome after elimination of redundancy and 0408 genome. The horizontal axis represents the GC content, and the vertical axis represents the Depth. Data were obtained in 10 Kb successively window.

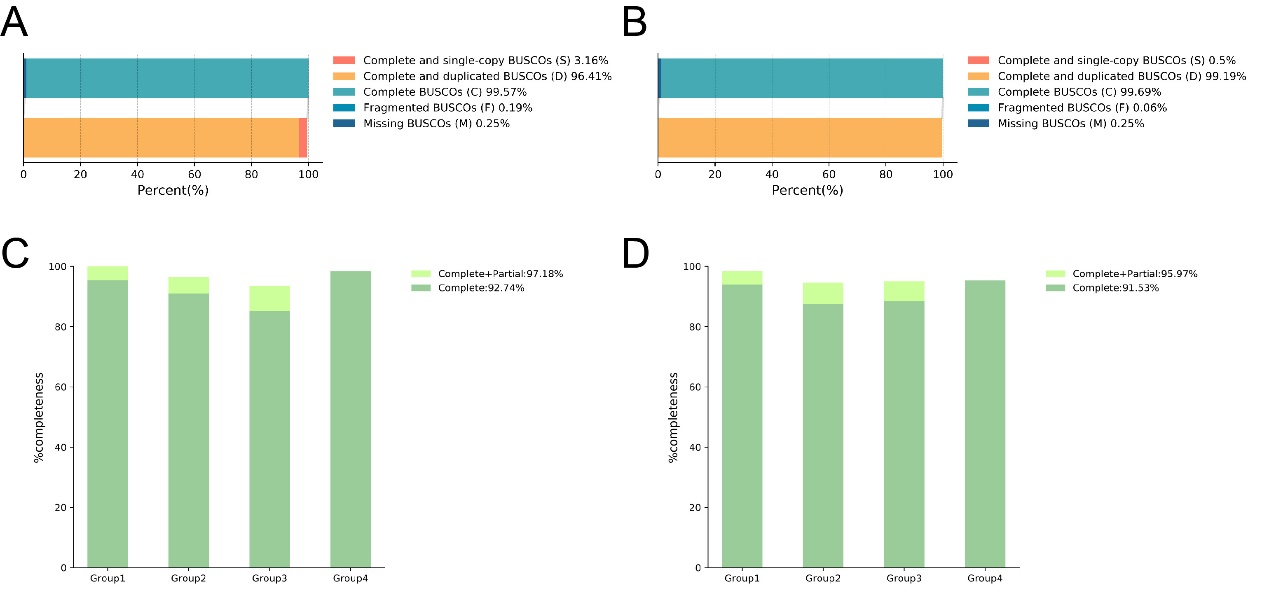

**Supplementary Figure S3.** Assessment of genome assembly completeness. (A-B) Genome assembly completeness was assessed using BUSCO for 0004 (A) and 0408 (B). (C-D) Conserved gene was assessed using CEGMA for 0004 (C) and 0408 (D).

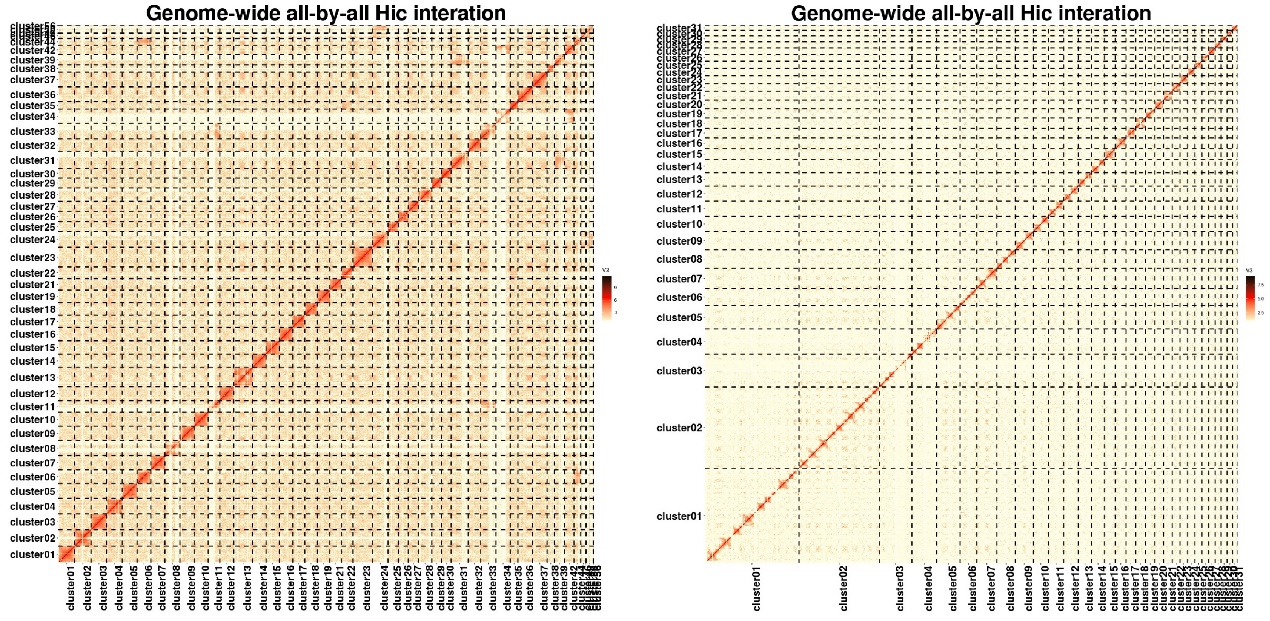

**Supplementary Figure S4.** Heatmap of primary Hi-C scaffolding of 0004 (left) and 0408 (right). The chromosome basic number was assumed to 56 for 0004 and 31 for 0408. The heatmap coordinates represent chromosomes, and the color of each point represents the log value of the corresponding genome bin pair interaction intensity, which increases sequentially from yellow to black.

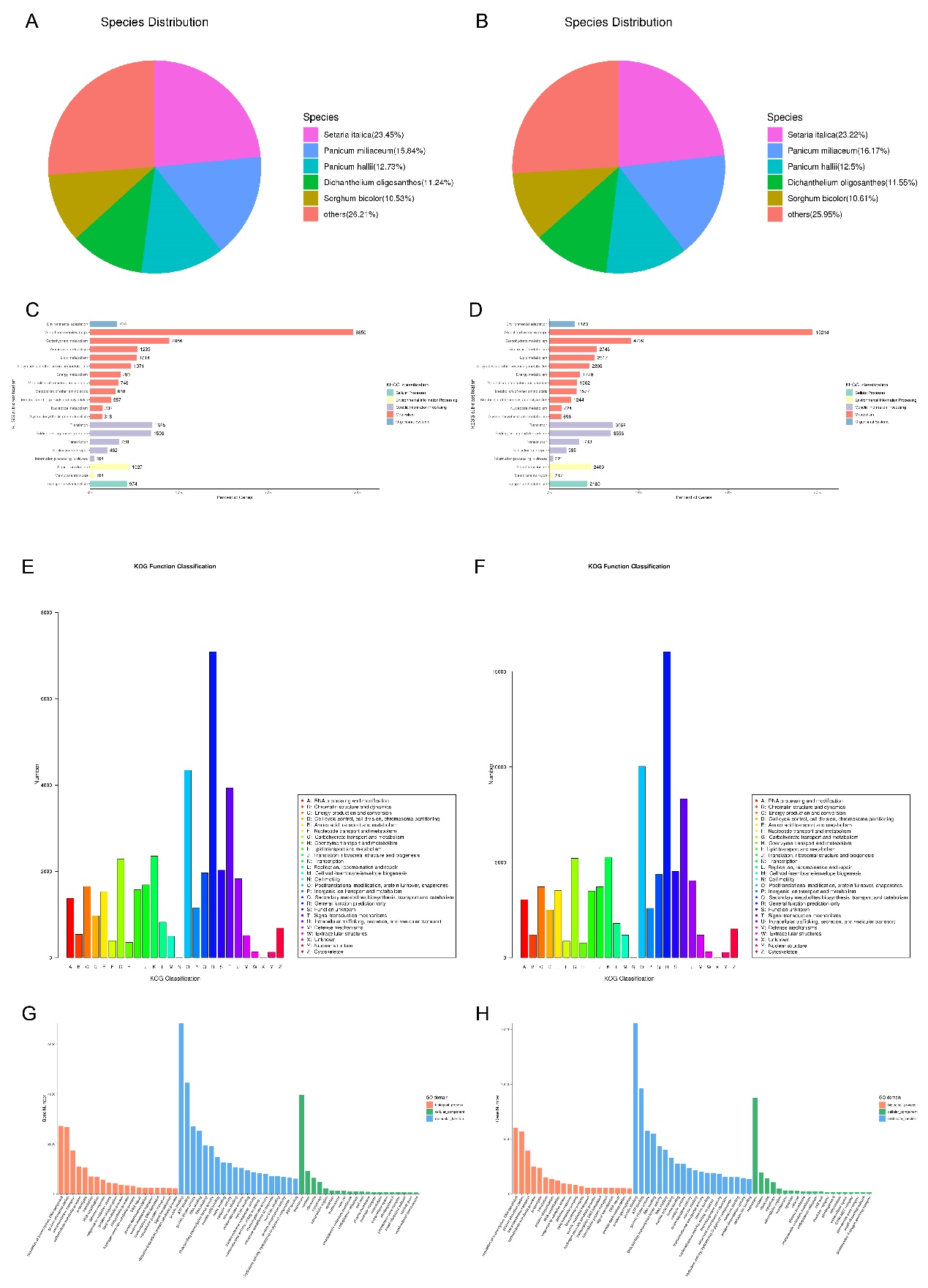

**Supplementary Figure S5.** Genome annotation based on public databases. (A-B) Species distribution for NR-based annotation for 0004 (A) and 0408 genome (B). (C-D) KEGG classification of 0004 (C) and 0408 genome (D) annotation. (E-F) KOG function classification of 0004 (E) and 0408 (F) genome annotation. (G-H) GO domain of 0004 (G) and 0408 (H) genome annotation.

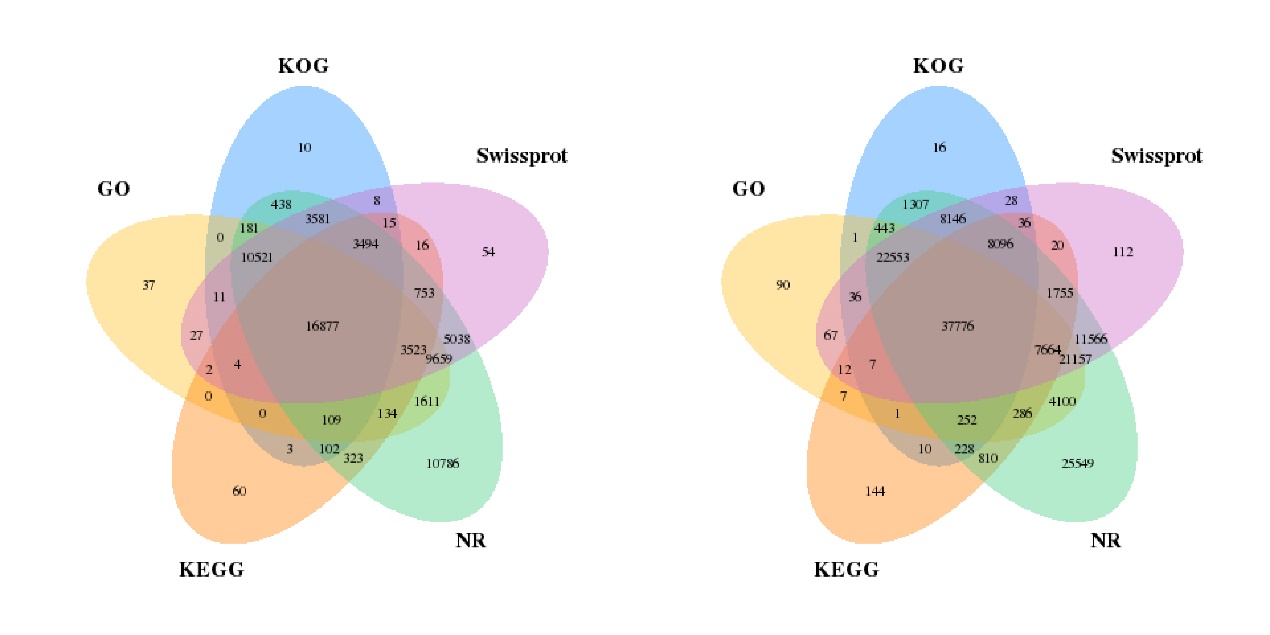

**Supplementary Figure S6.** Statistics of genome annotation for 0004 (left) and 0408 genome based on five databases, including Non-Reduntant Protein Database (NR), Kyoto Encyclopedia of Gene and Genomes (KEGG), Eukaryotic Orthologous Groups of protein (KOG), GO and Swissprot.

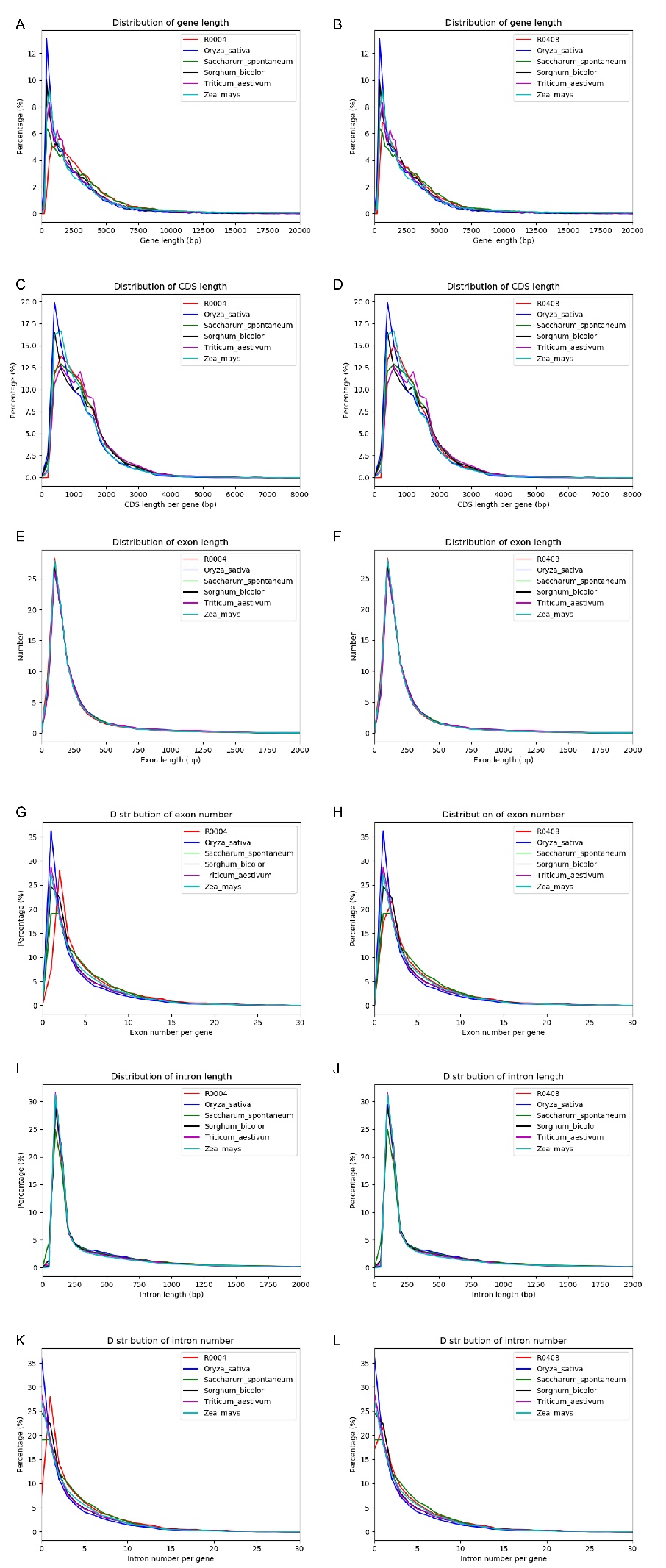

**Supplementary Figure S7.** Comparative statistics of gene information of related species. Comparison of gene information of 0004 (left column) and 0408 (right column) with five Poaceae plants (*Oryza sativa*, *Saccharum spontaneum*, *Sorghum bicolor*, *Zea mays* and *Triticum aestivum*) including distribution of gene length (A-B), CDS length (C-D), exon length (E-F), exon number (G-H), intron length (I-J) and intron number (K-L).

**Supplementary tables**

**Supplementary table 1** Statistics of RNA-seq data

| Sample name | Total reads | Total mapped | Unique mapped | Multiple  mapped |
| --- | --- | --- | --- | --- |
| 0004-root | 75,570,094 | 69,694,640 (92.23%) | 67,979,118 (89.96%) | 1,715,522 (2.27%) |
| 0004-leaf | 73,310,846 | 68,078,324 (92.86%) | 66,040,232 (90.08%) | 2,038,092 (2.78%) |
| 0004-panicle | 121,085,174 | 109,619,650 (90.53%) | 107,277,684 (88.60%) | 2,341,966 (1.93%) |
| 0004-bud | 102,532,136 | 95,570,548 (93.21%) | 93,230,426 (90.93%) | 2,340,122 (2.28%) |
| 0408-root | 112,487,910 | 110,342,744 (98.09%) | 95,485,032 (84.88%) | 14,857,712 (13.21%) |
| 0408-leaf | 81,606,088 | 79,939,116 (97.96%) | 68,384,754 (83.80%) | 11,554,362 (14.16%) |
| 0408-panicle | 62,558,910 | 59,462,320 (95.06%) | 50,570,542 (80.84%) | 8,891,778 (14.22%) |
| 0408-bud | 89,793,602 | 87,915,064 (97.91%) | 75,257,540 (83.81%) | 12,657,524 (14.10%) |
